# Inclusion of Oxford Nanopore long reads improves all microbial and phage metagenome-assembled genomes from a complex aquifer system

**DOI:** 10.1101/2019.12.18.880807

**Authors:** Will A. Overholt, Martin Hölzer, Patricia Geesink, Celia Diezel, Manja Marz, Kirsten Küsel

## Abstract

Assembling microbial and phage genomes from metagenomes is a powerful and appealing method to understand structure-function relationships in complex environments. In order to compare the recovery of genomes from microorganisms and their phages from groundwater, we generated shotgun metagenomes with Illumina sequencing accompanied by long reads derived from the Oxford Nanopore sequencing platform. Assembly and metagenome-assembled genome (MAG) metrics for both microbes and viruses were determined from Illumina-only assemblies and a hybrid assembly approach. Strikingly, the hybrid approach more than doubled the number of mid to high-quality MAGs (> 50% completion, < 10% redundancy), generated nearly four-fold more phage genomes, and improved all associated genome metrics relative to the Illumina only method. The hybrid assemblies yielded MAGs that were on average 7.8% more complete, with 133 fewer contigs and a 14 kbp greater N50. Furthermore, the longer contigs from the hybrid approach generated microbial MAGs that had a higher proportion of rRNA genes. We demonstrate this usefulness by linking microbial MAGs containing 16S rRNA genes with extensive amplicon dataset. This work provides quantitative data to inform a cost-benefit analysis on the decision to supplement shotgun metagenomic projects with long reads towards the goal of recovering genomes from environmentally abundant groups.

## Introduction

Shotgun metagenomics is a powerful method that conceptually allows all the genomes from all the organisms and their associated viruses within a sample to be determined with sufficient sequencing depth (Venter *et al.*, 2004; Handelsman *et al.*, 2007). In practice, metagenomic data typically represents hundreds to thousands of microorganisms and viruses at different coverage levels depending on the community structure within the sample (richness, evenness, and genome size variation). These data enable the determination of the community composition (who is there) and total community function (what are they capable of doing). In addition to this wealth of information, one of the most beneficial outcomes of shotgun metagenomic projects is the ability to assemble high quality, complete or nearly-complete, genomes from organisms not yet amenable to cultivation practices (Tyson *et al.*, 2004; Luo *et al.*, 2012). And indeed, such metagenome-assembled-genomes (MAGs) have provided information leading to the cultivation of organisms of interest (Gutleben *et al.*, 2018; Cross *et al.*, 2019; Imachi *et al.*, 2019), along with the discoveries of new metabolic processes (Daims *et al.*, 2015), novel insights into the ecology and evolution of globally abundant groups (Delmont *et al.*, 2018), and uncovering a wide diversity of novel Phylum-level lineages that have restructured the current understanding of the tree of life (Brown *et al.*, 2015; Hug *et al.*, 2016).

MAGs not only represent bacteria and archaea, but also include viruses, which are an integral part of most metagenomes (Dutilh *et al.*, 2014). Viruses are the most abundant biological entities in many ecosystems and can exert proportionately large effects on ecosystem functions (Fuhrman, 1999; Breitbart *et al.*, 2002). Viral MAGs have lead to the discovery of megaphages (with genomes >540 kb) from human and animal gut microbiomes (Devoto *et al.*, 2019), provided the first insights into the global distribution of such megaphages (Al-Shayeb *et al.*, 2019), and confirmed that environmental cyanophages contribute to global marine photosynthesis rates (Fridman *et al.*, 2017). However, the identification of viruses from metagenomes and their distinction from prophages continues to be challenging because there is not an established computational gold standard (Nooij *et al.*, 2018).

One exciting avenue in reconstructing *de novo* MAGs has been the inclusion of long-read sequences that can act as scaffolds to short-read sequences to help improve contiguity and bridge repeat regions within a genome (Chen *et al.*, 2019). This concept has been present since the advent of second-generation sequencing technologies (Roche 454, Illumina, SOLiD) (Goldberg *et al.*, 2006), but the breakthroughs in third-generation sequencing technologies, particularly PacBio and Oxford Nanopore Technologies (ONT), have improved the practicality of such approaches by providing access to much longer reads (Scholz *et al.*, 2014; Frank *et al.*, 2016; Bertrand *et al.*, 2019).

The goal of this study was to compare a hybrid assembly approach, incorporating ONT long reads, to an Illumina-only short-read approach with respect to the recovery of high-quality MAGs from a groundwater ecosystem. We leveraged the well-characterized monitoring transect of the Hanich Critical Zone Exploratory (CZE) that encompasses 15 monitoring wells spread across a hillslope covered by mixed beech forest, pasture land, and cropland (Küsel *et al.*, 2016). Groundwater from this site contains a wide diversity of microbial life. The microorganism component is dominated by Patescibacteria, uncultured organisms that are often missed by routine amplicon datasets (Herrmann *et al.*, 2019; Wegner *et al.*, 2019), and the viral component has only recently started to be explored (Kallies *et al.*, 2019). A previous, gene-centric based metagenomics project has identified dominant metabolic pathways within the aquifer (Wegner *et al.*, 2019). However, it has been challenging to directly link these key metabolic pathways to the specific microorganisms mediating them, which is critical for our understanding of the ecology of the site.

In order to address these knowledge gaps, we first quantified the improvements in the recovery of MAGs by including Oxford Nanopore Technology (ONT) long reads. This approach doubled the number of recovered MAGs that met our quality thresholds and covered a wider range of phylogenetic diversity present. The hybrid approach also improved all MAG metrics assessed and had a higher proportion of recovered MAGs with rRNA genes. From a viromics perspective, there were nearly four times more phages identified from the hybrid assembly, and of these, there were 10x more prophages identified. The results from this study are likely conservative and we expect there to be further improvements as ONT sequence quality increases with more accurate base calling algorithms and as more assembly and binning algorithms are developed to take advantage of all the information provided by long reads.

## Materials and Methods

### Sample Collection, DNA Extraction

Groundwater was collected in the Hainich Critical Zone Exploratory (NW Thuringia, Germany), from shallow groundwater resources in Upper Muschelkalk bedrock that has been extensively described (Küsel *et al.*, 2016; Lehmann and Totsche, 2020). In brief, the Hainich CZE contains a multistorey, fractured aquifer system within the hillslope, composed of altering layers of limestone and mudstone (Kohlhepp *et al.*, 2017). The Upper-Muschelkalk aquifer system is characterized by the limestone-dominated main aquifer (Trochitenkalk formation, moTK; formerly referred to as the HTL) that is predominantly oxic and the mudstone-dominated hanging strata (including Meissner formation, moM; formerly the HTU) that is anoxic (Kohlhepp *et al.*, 2017). Groundwater (115 L) was collected from well H52 (moM) on December 11th, 2018 and was sequentially filtered through 0.2 μM, and 0.1 μM PTFE filters (142 mm, Omnipore Membrane, Merck Millipore, Germany). Filters were immediately frozen on dry ice and transported to a −80° C freezer.

The DNA extraction was performed as previously described, using a phenol-chloroform based method without mechanical lysis to minimize fragmentation (Taubert *et al.*, 2018). Following extraction, the Zymo DNA Clean & Concentrator kit was used to purify and concentrate the DNA for both Illumina and Oxford Nanopore Technology (ONT) sequencing. DNA concentrations of 8.89 ng/μL (0.1μM filter fraction) and 37.7 ng/μL (0.2μM filter fraction) were measured using a Qubit 4 Fluorometer (Invitrogen).

### Illumina Metagenome Preparation and Initial Processing

Illumina libraries from both filter fractions were generated using the NEBNext Ultra II FS DNA library preparation kit following the recommended protocol. Size selection was performed using the AMPure XP beads (Beckman Coulter). The average insert sizes were 509 bp (0.2μM fraction) and 392 bp (0.1 μM fraction) as determined with an Agilent Bioanalyzer using a DNA7500 chip. The sequencing was performed in-house on an Illumina Miseq with 2×300 bp v3 chemistry. The 0.2 μM filter fraction DNA sample generated 17,660,385 paired-end sequences (10.6 Gbp), while the smaller 0.1 μM fraction sample generated 15,546,350 (9.33 Gbp) raw sequences.

Adapter sequences from the Illumina reads were removed using bbduk using kmer searching (k=23, hdist=1) and reads were trimmed with a phred score of 20, allowing to trim both sides of the read. Reads shorter than 50 bp were discarded (Bushnell, 2014). The average estimated insert sizes of the sequences were 234 bp and 214 for the 0.2 μM fraction and the 0.1 μM fraction samples, respectively. Raw reads were deposited at the ENA under accession PRJEB35315.

### Oxford Nanopore Metagenome Preparation

We performed Nanopore sequencing of the 0.2 μM fraction on a single MinION flow cell (FLO-MIN106 with an R9.4.1 pore) using the 1D genomic DNA by ligation kit (SQK-LSK109, ONT) following manufacturers’ instructions with minor adaptations. In short, the initial g-TUBE shearing step was omitted and potential nicks in DNA and DNA ends were repaired in a combined step using NEBNext FFPE DNA Repair Mix and NEBNext Ultra II End repair/dA-tailing Module (New England Biolabs, USA) and doubling the incubation time. A subsequent AMPure bead (Agencourt AMPure XP, Beckman Coulter) purification was followed by the ligation of sequencing adapters onto prepared ends. A second clean-up step with AMPure beads was performed and sequencing buffer and loading beads were added to the library. An initial quality check of the flow cell (ID: FAK43462) showed 1761 active pores at the start of sequencing. We loaded the DNA with a concentration of 98 ng/ul (measured by Qubit 3 Fluorometer; Thermo Fisher Scientific) and a total amount of ~1.4 μg. The sequencing run stalled after 18 h and was restarted for another 24 h using the MinKNOW software.

Basecalling was performed using the Guppy software (v2.3.1) with the high-accuracy model r9.4.1_450bps_large_flipflop. Called reads were classified as either pass or fail depending on their mean quality score. A total of 2,380,279 reads were basecalled and of these 2,081,879 (87.5%) were passed as satisfying the quality metric. The passed reads contain a total of 11.58 Gb of DNA sequence with a mean read length of 5,560 nt. This passed-fraction amounts to 91.8% of the total DNA nucleotide bases sequenced.

We deposited the raw signal files (FAST5) and basecalled reads (FASTQ) at the ENA under accession PRJEB35315.

### Metagenome Assembly and Binning

We used metaSPAdes v3.13.0 (Nurk *et al.*, 2017) to assemble the hybrid and Illumina-only data in order to be able to directly compare both approaches. The sequences from the 0.2 μm sample were individually assembled with the default parameters specified by the “-meta” flag with only the inclusion of the long reads with the “-nanopore” flag being different. The assembly statistics for the two methods were calculated using MetaQUAST with the default settings (Mikheenko *et al.*, 2016). We binned the scaffolds from each assembly that were longer than 1000 nucleotides (nt) using MaxBin2 (Wu *et al.*, 2016) and MetaBAT2 (Kang *et al.*, 2015, 2019) included in the MetaWRAP (Uritskiy *et al.*, 2018) binning module with the “--universal” flag. Differential coverage information was included using Illumina QA/QC reads from both filter fractions. Additionally, scaffolds > 3000 nt from each assembly were binned with BinSanity using the “wf” workflow and the log normalized coverage file produced with the “BinSanity-profile” command (Graham *et al.*, 2017). The MetaWRAP “Bin_refinement” module was used to dereplicate bins produced from the 3 different binning methods, using the MetaWRAP scoring algorithm which favors low redundancy values while also selecting for higher percent completion. We found that the MetaWRAP “Ressamble_bins” module reduced the quality of our bins and thus proceeded with the two sets of refined bins that were at least 50% complete with less than 10% redundancy.

Each collection was imported into Anvi’o and MAG statistics were exported using the anvi-summarize command (Murat Eren *et al.*, 2015). Our completeness and redundancy/contamination criteria were initially assessed using estimations from checkM (Parks *et al.*, 2015), while the estimations exported from anvio were used to calculate the values represented in Figure 1 (visualized using the R package ggplot2 v3.1.0 (Wickham, 2009; Wickham *et al.*, 2019)). The anvio completeness estimations were lower than the checkM values while the redundancy/contamination estimations were higher, and therefore 74 hybrid and 39 Illumina-only MAGs were included, out of the original 82 and 44. Each MAG was screened for rRNA genes using barrnap, with each hit required to be at least 20% of the full gene length (Seemann, 2015). These statistics were compiled from the automatically refined MAGs and not from manually curated MAGs as we considered this the most direct comparison. While we consider it essential that MAGs be manually curated before publishing (Bowers *et al.*, 2017; Shaiber and Eren, 2019), using the automated results for this specific comparison minimizes added bias, and likely underestimates the actual improvements due to the more fragmentary nature of the Illumina-only MAGs.

**Figure 1.**
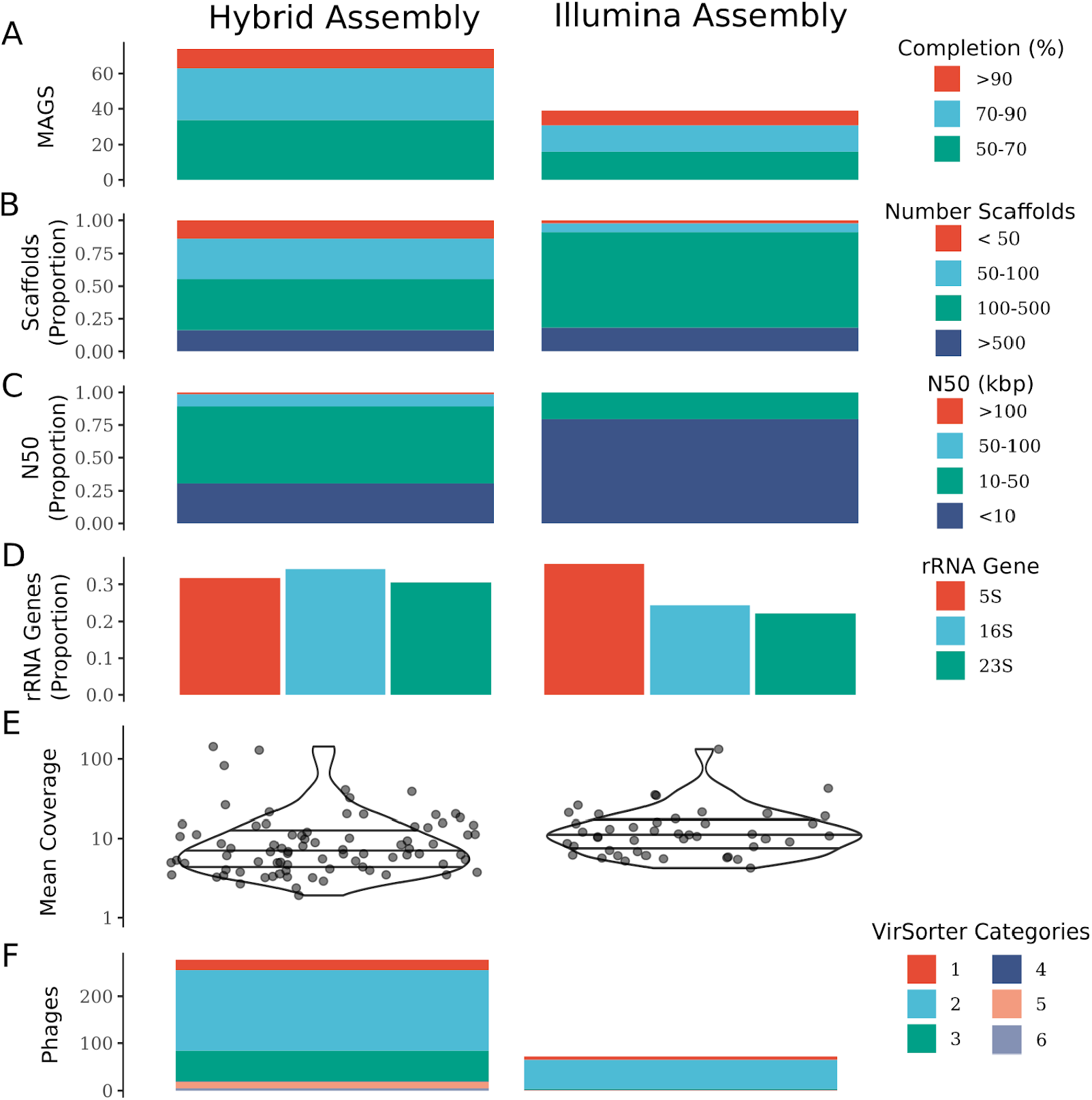
Genome metrics for all MAGs that were > 50% complete and < 10% redundant reconstructed from the hybrid assembly (left) and the Illumina-only assembly (right). (A) The number of MAGs from each method. (B) The number of scaffolds each MAG was represented by relative to all recovered MAGs. (C) The N50 value for each MAG recovered as a proportion of all MAGs. (D) The proportion of MAGs containing each of the rRNA genes. (E) The mean coverage of every MAG that met our quality standards within each approach (points) and their distributions plotted on a log10 scale. (F) The number of identified phages / prophages in each assembly as assessed with VirSorter.

To identify MAGs that were recovered from both assembly methods, FastANI was run on all pairwise comparisons of MAGs that were initially assessed using checkM (82 hybrid, 44 Illumina-only) (Jain *et al.*, 2018). All MAGs that were recovered from both assemblies had an average nucleotide identity (ANI) of at least 98.8%, and in all cases, secondary hits were < 82% ANI. These shared MAGs were investigated in more detail as they enabled a direct comparison between the Illumina and the hybrid assemblies. Paired Welch’s t-tests with Benjamini-Hochberg correction were used to specifically test the differences in completeness, redundancy/contamination, genome length, number of scaffolds, and N50. The proportion of MAGs containing each of the rRNA genes was tested using a two-sample z-test in R (R Core Team, 2014).

The scaffolds from each pair of MAGs that were recovered were aligned using mummer wrapped within the mummer2circos.py (https://github.com/metagenlab/mummer2circos) script (Kurtz *et al.*, 2004). All nanopore reads > 1 kbp were aligned to a concatenated fasta file of all scaffolds from all the hybrid-generated bins using minimap2 with the “-ax map-ont” flag (Li, 2018). Log2 scaled coverage profiles were generated from QAQC Illumina reads for each filter fraction using pileup.sh across 1 kb sized sections from the BBTools suite (Bushnell, 2014), after aligning the reads to the hybrid-bin scaffolds using bbmap.sh and sorting the alignments with samtools (Li *et al.*, 2009). MAGs were taxonomically classified using GTDB-TK v0.3.2, following the classify workflow (Hyatt *et al.*, 2010; Matsen *et al.*, 2010; Price *et al.*, 2010; Eddy, 2017; Jain *et al.*, 2018; Parks *et al.*, 2018).

The 16S rRNA genes recovered from MAGs were compared using blastn to 97% representative operational taxonomic units (OTUs) from primer pair 341F/785R (Altschul *et al.*, 1990; Yan *et al.*, 2019). All hits were required to be >98.5 % sequence identity across at least 350 bp. In the case of multiple OTUs matching a MAG 16S gene, the highest bit score was used, followed by the most abundant representative OTU. The representative OTUs originated from 101 samples collected between July 2014 and April 2017 from 10 monitoring wells of the hillslope transect (Yan *et al.*, 2019). The figure was created using ggplot2 and the tidyverse package within R (R Core Team, 2014; Wickham *et al.*, 2019).

In order to disentangle the improvements in MAG recovery due to longer ONT scaffolds instead of simply having higher sequencing depth, we subsampled the ONT reads into four different sets, then re-ran the SPAdes assemblies and binning steps. As described above, the full hybrid assembly approach was based on 17,660,385 Illumina paired-end sequences (10.6 Gbp), and 2,081,879 ONT sequences (11.58 Gbp). The four additional ONT sets were, (1) all nanopore reads >10,000 nt which lead to 349,321 sequences (6.8 Gbp), (2) ONT reads > 20,000 nt (114,853 sequences; 3.6 Gbp), (3) ONT reads > 50,000 nt (7,848 sequences; 0.48 Gbp), and (4) randomly subsampled to 25% of the initial number of sequences (595,070 sequences, 3.15 Gbp). Each of the resulting assemblies were binned and analyzed following the same procedures as above.

### Viral Comparison

We used VirSorter v1.0.5 to search for putative phage and prophage sequences in the assemblies (Roux *et al.*, 2015). To identify the phages that were recovered from both assemblies, we used Blastn v2.9.0+ and filtered the hits by an e-value of 1e-10, a sequence identity >90%, and an alignment length >50% (Altschul *et al.*, 1990).

## Results and Discussion

One of the most striking findings is that the inclusion of long reads more than doubled (74 vs 39) the number of bacterial and archaeal MAGs that were at least 50% complete and less than 10% redundant, as compared to using assemblies generated from only Illumina short-read sequences (Figure 1 A). The 74 MAGs recovered from the hybrid approach were less fragmented with more than 50% having less than 100 scaffolds as compared to ~5% of the Illumina-only MAGs (Figure 1 B). The hybrid MAGs correspondingly exhibited much higher N50 values with > 75% of the MAGs having an N50 > 10 kbp as compared to 20% of the Illumina-only MAGs (Figure 1 C). The additional MAGs recovered from the hybrid approach were due to recovering MAGs with lower coverages (minimum 1.9x vs 4.2x) and improved recovery of populations that were close to our quality cutoff values (Figure 1 E).

In addition to the bacterial and archaeal MAGs, we identified 278 putative phage sequences (258 phages, 20 prophages) in the hybrid assembly and only 73 (71 phages, 2 prophages) in the Illumina-only assembly (Figure 1 F). Remarkably, the number of complete phage contigs of the VirSorter category 1 (confident phage assignment) increased from 7 to 22 sequences in the hybrid assembly. Thus, the integration of long-read data into the assembly process helped to recover more potential phage sequences from the sample, as well as an increase in phage sequences that were confidently identified. In one noteworthy case, a category 2 phage sequence (enriched in viral domains) from the Illumina-only assembly (length 17,648 nt) was integrated into a much larger prophage contig (category 4; high confidence prophage) within the hybrid assembly (75,806 nt). Of the 73 identified putative phage sequences from the Illumina-only assembly, 52 were also recovered with the hybrid approach. The remaining 21 phage sequences are composed of three category-1, 17 category-2, and a single category-4 prophage. Of those, 19 sequences could be confirmed by an additional blast step to be included in larger contigs of the full hybrid assembly, however, they were not identified by VirSorter within the hybrid assembly.

For the bacteria and archaea, every MAG reconstructed from the Illumina-only assembly was also recovered from the hybrid assembly. We performed an in-depth comparison of the 44 bacteria and archaea MAGs that were initially recovered with both assemblies (Table S1). On average, the hybrid-assembled MAGs were 7.8% ± 3.1% (mean ± 95% confidence intervals) more complete (adj.p < 1e-5) and exhibited the same degree of redundancy (0.4% ± 0.7%; adj.p = 0.22). In addition, the hybrid MAGs were 358 ± 123 kbp longer (adj.p < 1e-5), with 133 ± 34 fewer scaffolds (adj.p < 1e-8), that had 14.1 ± 5.4 kbp greater N50 values (adj.p < 1e-5). There was no significant difference found in the proportion of the rRNA genes between the recovered MAGs (two proportion two-tailed z-test), although the hybrid MAGs had 1 more 5S gene (16 vs. 15), 6 more 16S genes (16 vs. 10), and 4 more 23S genes (13 vs. 9). A non-significant result is not discouraging here as even a single extra 16S rRNA gene might allow a dominant population to be connected to an extensive amplicon dataset. In addition, a manual examination of the few Illumina-only MAGs that were more complete than the corresponding hybrid MAG found that the Illumina-only MAGs were likely mixed populations with different coverage profiles across their scaffolds.

To see if the improvements in MAG-retrieval were mainly due to the increase in sequencing depth provided by the ONT reads, we subsampled using four different strategies. These ranged from only using ONT reads > 50,000 nt (7,848 sequences; 0.48 Gbp) to all reads > 10,000 nt (349,321; 6.8 Gbp), along with a random subsampling to simulate a less successful sequencing run (595,070 sequences; 3.15 Gbp) (Table S2). The randomly subsampled ONT set yielded 62 MAGs that were > 50% complete and < 10% redundancy (as assessed by checkM), 32 more MAGs than the MiSeq assembly alone. Including only the ~8,000 reads that were greater than 50,000 nt resulted in 58 MAGs passing our standards. We interpret these results to show that the longer nanopore reads do help recover MAGs, and a large improvement can be expected even if the ONT run was not so successful, while also acknowledging that the greater sequencing depth alone supplied by the ONT reads partially contributes to these results.

The full hybrid assembly recovered MAGs representing 17 phyla, while 14 phyla were represented from the Illumina-only assembly. The missing phyla were Bacteroidota (2 MAGs), Microarchaeota (1), and Verrucomicrobiota (1). The (super)phylum with the greatest improvement in the number of assembled MAGs was the Patescibacteria (formerly the Candidate Phyla Radiation group), with over double (35 vs. 17) the number of MAGs recovered from the hybrid approach that met our criteria (Figure 2). The ONT long-read sequences bridge missing gaps in the Illumina-only MAGs, thereby improving the contiguity and increasing the genome length across fewer scaffolds (Figure 2). It is noteworthy that these benefits are not restricted to only the most abundant organisms and even relatively few long-reads mapping to populations can improve chances to recover them (Figure 2 B). Here we focused on the Patescibacteria, since this group contains almost no cultivated representatives and they are often highly abundant in groundwater systems (Brown *et al.*, 2015; Pedron *et al.*, 2019). In addition, previous research from the Hainch CZE has demonstrated that Patescibacteria can represent up to 79% of the groundwater community (Herrmann *et al.*, 2019).

**Figure 2.**
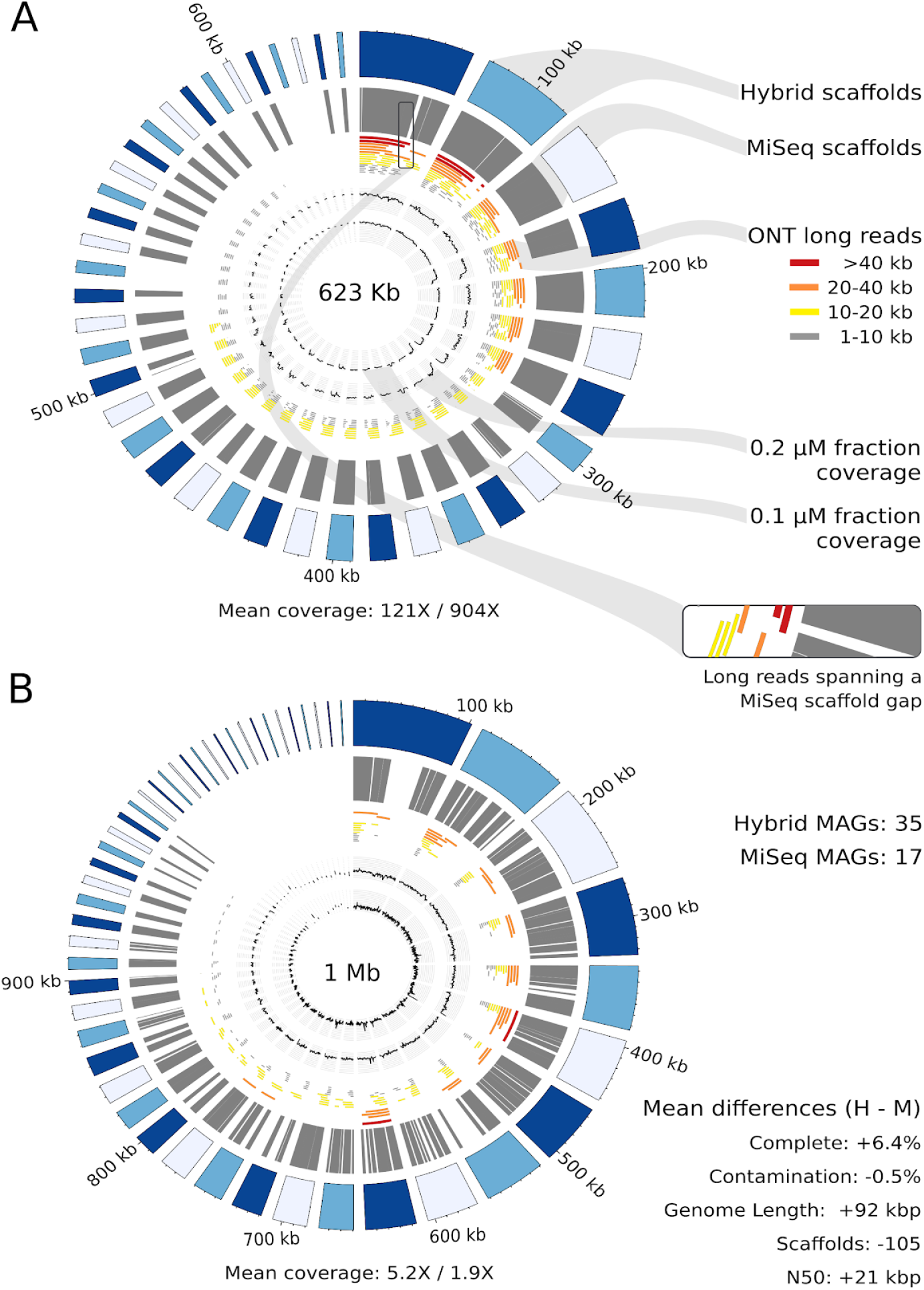
Genome circos plots for the most (A) and least (B) covered Patescibacteria MAGs retrieved by both assembly methods. The outer ring in blue represents the hybrid assembly derived scaffolds, followed by the corresponding Illumina assembly scaffolds in grey. The nanopore reads were mapped with minimap2 and colored based on length. The coverage values are log2 scaled and calculated for each 1kb segment of the hybrid-derived scaffolds with pileup.sh from BBTools of the Illumina reads. The values below each plot represent the mean coverage from the 0.2 μM fraction Illumina MiSeq reads and the 0.1 μM fraction reads, respectively. The hybrid based genome size is indicated in the middle of each plot.

A similar visualization was performed for three other dominant bacterial phyla (Nitrospirota, Actinobacteriota, Proteobacteria) that had previously been shown to be important in the functioning of the groundwater (Wegner *et al.*, 2019; Yan *et al.*, 2019) (Figure S1). There were almost double the number of Nitrospirota associated MAGs (7 vs. 4) recovered with the hybrid approach, and these were amongst the largest of the MAGs recovered. Within the Actinobacteriota and Proteobacteria, we recovered the same MAGs with each approach. The MAG metrics within each phyla fell within the confidence intervals of the full sample set with the exception of the Actinobacteriota that showed greater than expected improvements with the hybrid approach (Figure S1). The results presented herein also likely further underestimate the improvements ONT long reads contribute since the 2×300 bp paired-end sequences used are longer than those typically generated in metagenome projects (either 2×150 reads from an Illumina HiSeq or 2×250 from an Illumina Nextseq).

Our results from these diverse and understudied groundwater ecosystems extend findings recently published that utilized mock communities and spiked-in complex human gut microbiome communities (Bertrand *et al.*, 2019). We demonstrate the recovery of a wider diversity of microorganisms and phages using a hybrid approach, in our case, the recovery of three additional phyla and 6 classes. Additionally, the improvements towards contiguity and completeness of recovered MAGs are likely generalizable as they are reflected within the results presented by Bertrand and colleagues (2019). While not explicitly tested in our study, the hybrid approach has been previously shown to result in fewer misassemblies using a mock community where the genome contents are known *a priori* (Bertrand *et al.*, 2019).

High-quality MAGs that are (nearly) complete with lower frequencies of misassemblies allow relevant metabolic processes to be constrained to individual organisms, directly connecting phylogeny to function (Woyke *et al.*, 2019). To help with this task, one of the benefits we document here is that the less fragmented hybrid MAGs contained more ribosomal RNA genes (Figure 1 D). 16S rRNA gene amplicon datasets are a common method to survey microbial communities, often with a large number of samples that are well replicated spatiotemporally due to decreases in sequencing costs and ease of multiplexing. Here, MAGs were linked to a 16S dataset containing 101 samples from the Hainich CZE that were collected across 3 years (July 2014 to April 2017) and 10 wells (Yan *et al.*, 2019).

Of the 16S hybrid MAGs that mapped to a 16S OTU, four were visualized based on their relative abundance and spatiotemporal distributions throughout the Hainich CZE (Figure 3). The 16S sequence from bin49 was full length (1498 bp) and shared 100% identity across the full OTU_31 sequence. The MAG taxonomy (GTDB_TK) was class ABY1 within the Patescibacteria, while the 16S taxonomy (RDP trained with SILVA v.132) was *Candidatus Kerfeldbacteria* within the same class (Wang *et al.*, 2007; Schloss *et al.*, 2009; Quast *et al.*, 2012; Parks *et al.*, 2018). This population recently showed a large increase in relative abundance starting in Jan 2017 and is localized to well H5-2 within the upper aquifer assemblage, where these metagenomes originated from. Bin19 and bin52 also both contained partial 16S sequences (769 bp, 727 bp respectively) that were 100% identical to corresponding 16S OTU sequences. The GTDB_TK taxonomy for bin19 was within the order Peregrinibacterales (Patescibacteria) while bin52 was most similar to the MBNT15 group. One exciting finding was that both these organisms were more relatively abundant in the oxic well H4-1 within the lower aquifer assemblage rather than the anoxic well that was used to generate the metagenomes. The last example provided is for a MAG, bin61, within the Thermodesulfovibrionales (Nitrospirota) that shared 99% sequence identity to the most abundant OTU detected across the entire Hainich CZE (mean ± SD relative abundance = 5.9 ± 7.6%; max = 32%). This OTU is extremely abundant within the anoxic wells H53 and H52, while also being among the most abundant OTU in the oxic or hypoxic Trochitenkalk formation (moTK; HTL) at times (wells H41 and H51). With these examples, we extend the information provided by a spatially constrained metagenomics project across the entire aquifer assemblage to better explore the potential niches of these abundant organisms.

**Figure 3.**
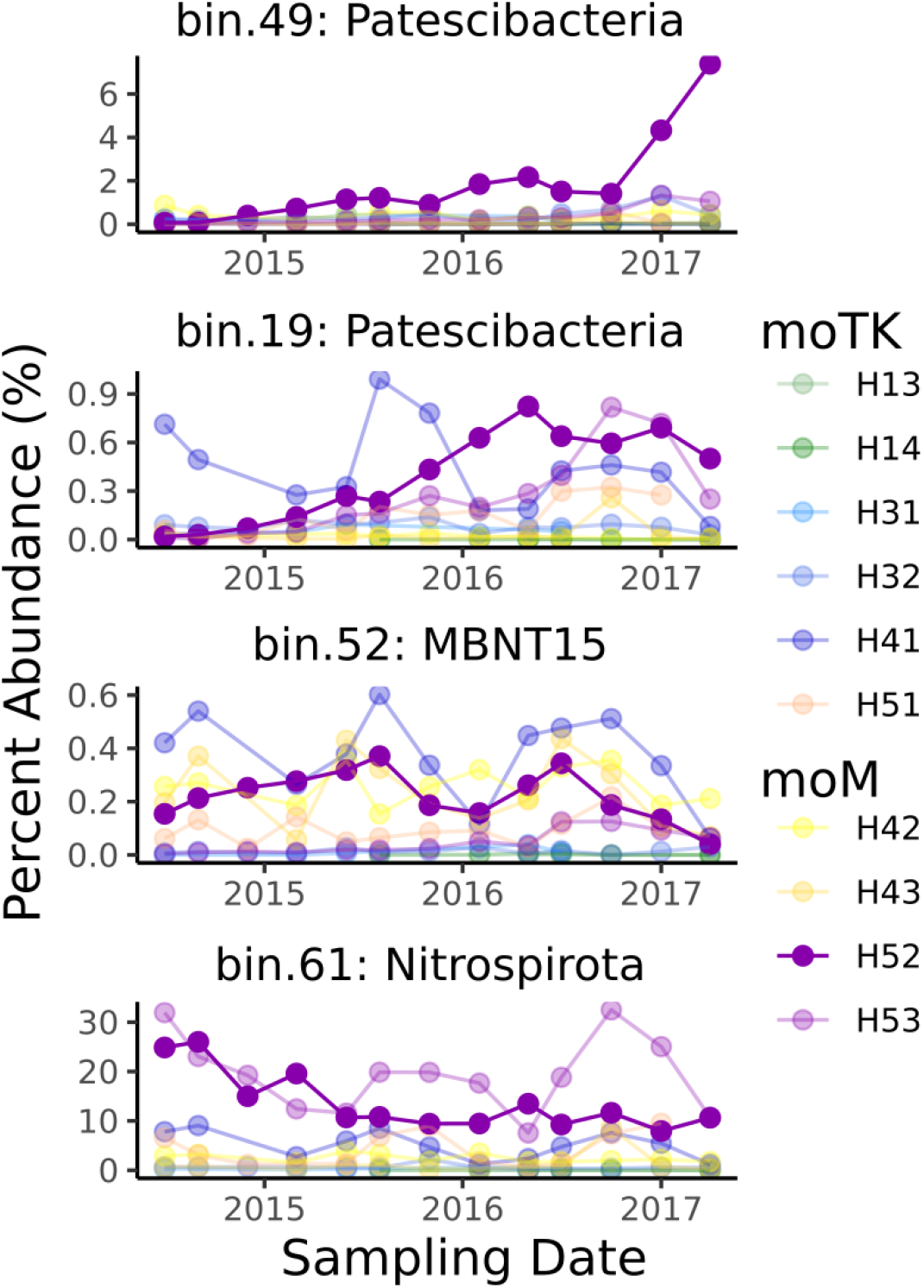
MAGs containing 16S genes were mapped to the amplicon sequence dataset presented in Yan et al., using blastn. All matches were > 99% sequence identity across at least 350 bp. The distribution of these OTU was explored throughout the full aquifer system across a 3-year monthly sampling time series. The aquifer is mainly divided into two assemblages, the upper (moM; HTU) characterized by anoxia, and the lower (moTK; HTL) by oxic and hypoxic conditions. H52 was the well where the DNA for the metagenomes originated from and is highlighted with the solid purple. The percent abundance values result from counts that were normalized with metagenomeseq (Paulson *et al.*, 2013). The mapping information is as follows: bin.49 (len = 1498 bp; ID = 100%, 404 bp); bin.19 (len = 769, ID = 100%, 405 bp); bin.52 (len = 769, ID = 100%, 405 bp); bin.61 (len = 1403; 99%, 402 bp). All 16S and MAG taxonomic assignments were consistent.

Unlike more traditional, gene-centric based analyzes that provide insights into the sum metabolic repertoire of an ecosystem, a genome-centric approach enables research questions directed towards population niche differentiation, determination of microbial groups that are bioindicators for a specific metabolism, and potential microbial networks and microbial interactions between syntrophs and/or auxotrophs along with the phages that control population sizes and alter or enhance biogeochemical cycling rates (Anantharaman *et al.*, 2016; Howard-Varona *et al.*, 2017). Such an approach further offers insights into the evolutionary history of archaeal, bacterial, or viral groups and the ecological consequences if such groups were to be lost or invade the system. In particular, the identification of novel viruses from metagenomic data and the way they interact with other microbes extends our understanding of complex environmental systems (Roux *et al.*, 2016). As a final consideration, the information contained within high-quality MAGs may offer a road map to cultivation, which in turn allows hypothesis testing and verification of *in silico* predictions (Cross *et al.*, 2019).

## Conclusions

To improve the recovery of metagenome-assembled-genomes we find that the addition of Oxford Nanopore Technology (ONT) long-read sequencing doubled the number of bacterial and archaeal MAGs, that represented more phylogenetic diversity, and improved all measured quality metrics as compared to an Illumina short-read approach only. In addition, nearly four-fold more putative phage sequences were identified including 10x more putative prophages. Considerations on supplementing Illumina paired-end metagenomic projects with ONT reads include the DNA extraction method used, the total amount of DNA available, and the cleanliness of the extract. The additional amount of hands-on time needed to prepare a sequencing run with the minION is comparably low. The current library preparation time using the revised genomic DNA by ligation kit takes about three hours (including elongated incubation times). Shorter protocols such as the rapid kit are also available by the ONT-community. A sequencing run lasts between 24 and 48 hours, or until no active pores are available anymore and the data can be immediately analyzed depending on the available hardware. There are cost concerns with supplementing an already expensive metagenomic sequencing project with ONT long-read sequences, considering a complete run on one minION flow cell, including library preparation, currently costs 750 € per sample. However, the improvements documented here provide better genome context for both microbial and viral comparative genomic projects, more single marker copy genes for detailed phylogenomic studies, more complete metabolic reconstructions, and an end-product that is more useful to the greater scientific community.

## Acknowledgements

We thank Falko Gutmann, Perla Abigail Figueroa-Gonzalez, Till Bornemann and Heiko Minkmar for water sampling and water filtration. We would also like to thank Alexander Probst for his helpful comments and discussion, and Stefan Riedel for operating the Illumina MiSeq. Martin Hölzer appreciates the support of the Joachim Herz Foundation by the add-on fellowship for interdisciplinary life science. This study was conducted within the Collaborative Research Centre AquaDiva (CRC 1076 AquaDiva) of the Friedrich Schiller University Jena, and was funded by the Deutsche Forschungsgemeinschaft.

The authors have no conflicts of interest.

---

**Figure S1.**
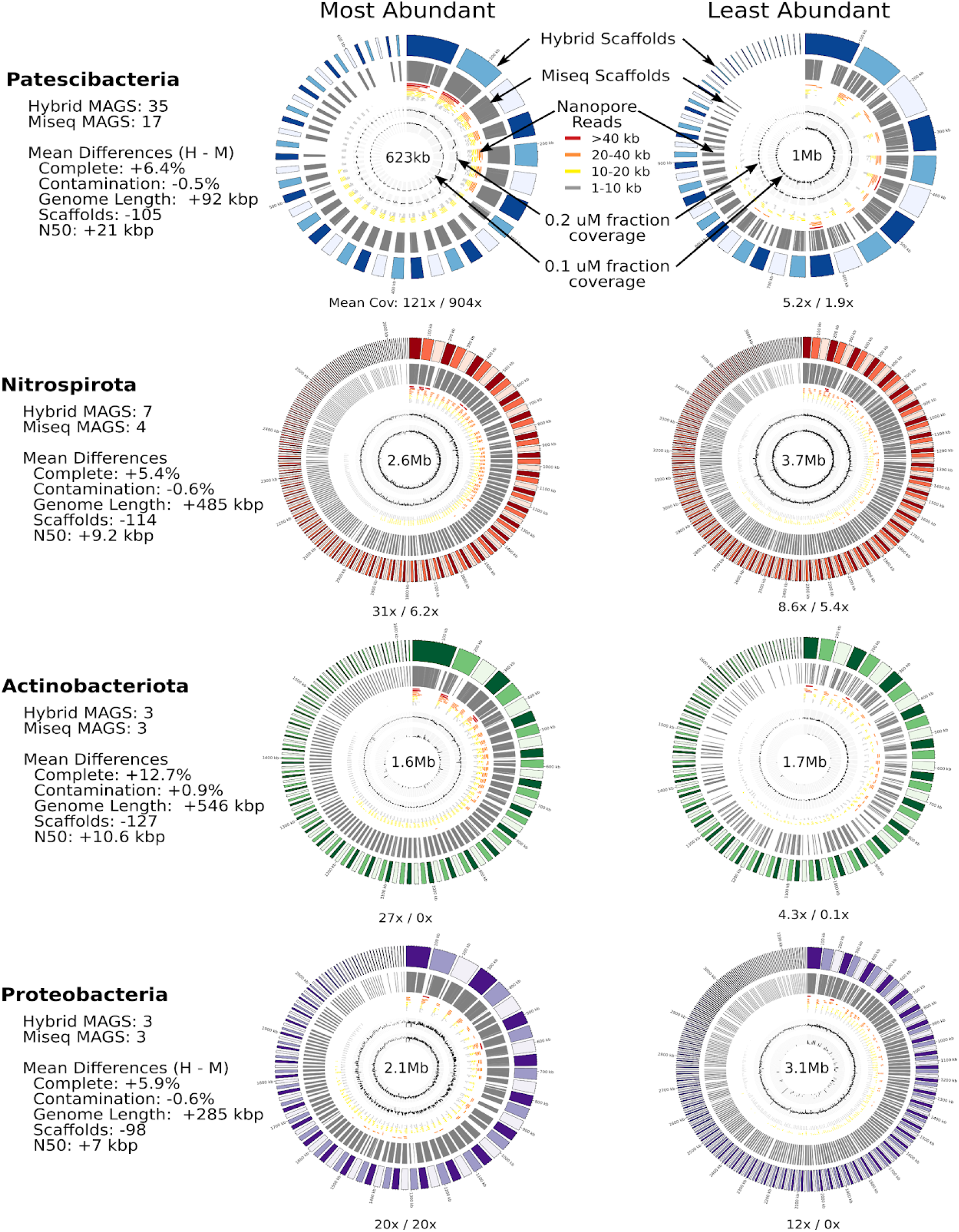
Genome circos plots for the most (left) and least (right) covered MAGs from four phyla previously demonstrated to be abundant and important in biogeochemical cycling in the Hainich CZE. The outer ring of each plot represents the hybrid assembly derived scaffolds, followed by the corresponding Illumina assembly scaffolds in grey. The ONT long reads were mapped with minimap2 and colored based on length. The coverage values are log2 scaled and calculated for each 1kb segment of the hybrid-derived scaffolds with pileup.sh from BBTools. The values below each plot represent the mean coverage from the 0.2 μM fraction Illumina MiSeq reads and the 0.1 μM fraction reads, respectively. The hybrid based genome size is indicated in the middle of each plot.

Table S1. Comparison of the MAGs (bins) that were recovered in both the hydrid and Illumina-only (Miseq) based analyses.

https://docs.google.com/spreadsheets/d/1AsoGrB6wfRGWqE6ToRQoHR8AWJq97A16_5UP-717c_0/edit#gid=2054870087

**Table S2.**
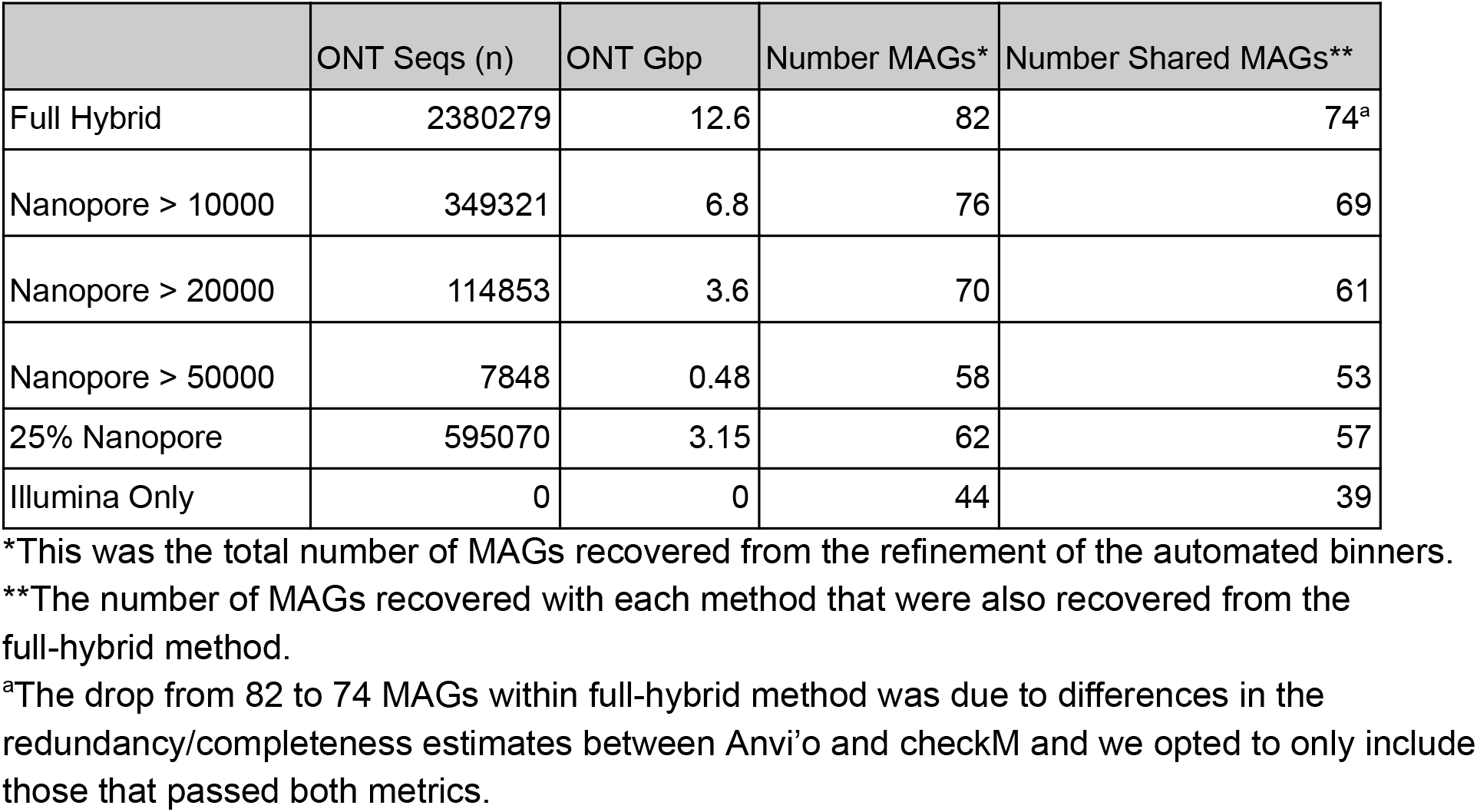
The effect of subsampling the ONT reads on the number of MAGs that were > 50% complete and < 10% redundant (as assessed by checkM).

## References

Al-Shayeb, B., Sachdeva, R., Chen, L.-X., Ward, F., Munk, P., Devoto, A., et al. (2019) Clades of huge phage from across Earth’s ecosystems. bioRxiv 572362.

Altschul, S.F., Gish, W., Miller, W., Myers, E.W., and Lipman, D.J. (1990) Basic local alignment search tool. J Mol Biol 215: 403–410.

Anantharaman, K., Brown, C.T., Hug, L.A., Sharon, I., Castelle, C.J., Probst, A.J., et al. (2016) Thousands of microbial genomes shed light on interconnected biogeochemical processes in an aquifer system. Nat Commun 7: 13219.

Bertrand, D., Shaw, J., Kalathiyappan, M., Ng, A.H.Q., Kumar, M.S., Li, C., et al. (2019) Hybrid metagenomic assembly enables high-resolution analysis of resistance determinants and mobile elements in human microbiomes. Nat Biotechnol 37: 937–944.

Bowers, R.M., Kyrpides, N.C., Stepanauskas, R., Harmon-Smith, M., Doud, D., Reddy, T.B.K., et al. (2017) Minimum information about a single amplified genome (MISAG) and a metagenome-assembled genome (MIMAG) of bacteria and archaea. Nat Biotechnol 35: 725–731.

Breitbart, M., Salamon, P., Andresen, B., Mahaffy, J.M., Segall, A.M., Mead, D., et al. (2002) Genomic analysis of uncultured marine viral communities. Proc Natl Acad Sci U S A 99: 14250–14255.

Brown, C.T., Hug, L.A., Thomas, B.C., Sharon, I., Castelle, C.J., Singh, A., et al. (2015) Unusual biology across a group comprising more than 15% of domain Bacteria. Nature 523: 208–211.

Bushnell, B. (2014) BBTools software package. URL http://sourceforgenet/projects/bbmap.

Chen, L.-X., Anantharaman, K., Shaiber, A., Murat Eren, A., and Banfield, J.F. (2019) Accurate and Complete Genomes from Metagenomes. bioRxiv 808410.

Cross, K.L., Campbell, J.H., Balachandran, M., Campbell, A.G., Cooper, S.J., Griffen, A., et al. (2019) Targeted isolation and cultivation of uncultivated bacteria by reverse genomics. Nat Biotechnol.

Daims, H., Lebedeva, E.V., Pjevac, P., Han, P., Herbold, C., Albertsen, M., et al. (2015) Complete nitrification by Nitrospira bacteria. Nature 528: 504–509.

Delmont, T.O., Quince, C., Shaiber, A., Esen, Ö.C., Lee, S.T., Rappé, M.S., et al. (2018) Nitrogen-fixing populations of Planctomycetes and Proteobacteria are abundant in surface ocean metagenomes. Nat Microbiol 3: 804–813.

Devoto, A.E., Santini, J.M., Olm, M.R., Anantharaman, K., Munk, P., Tung, J., et al. (2019) Megaphages infect Prevotella and variants are widespread in gut microbiomes. Nat Microbiol 4: 693–700.

Dutilh, B.E., Cassman, N., McNair, K., Sanchez, S.E., Silva, G.G.Z., Boling, L., et al. (2014) A highly abundant bacteriophage discovered in the unknown sequences of human faecal metagenomes. Nat Commun 5: 4498.

Eddy, S. (2017) HMMER3: a new generation of sequence homology search software.

Frank, J.A., Pan, Y., Tooming-Klunderud, A., Eijsink, V.G.H., McHardy, A.C., Nederbragt, A.J., and Pope, P.B. (2016) Improved metagenome assemblies and taxonomic binning using long-read circular consensus sequence data. Sci Rep 6: 25373.

Fridman, S., Flores-Uribe, J., Larom, S., Alalouf, O., Liran, O., Yacoby, I., et al. (2017) A myovirus encoding both photosystem I and II proteins enhances cyclic electron flow in infected Prochlorococcus cells. Nat Microbiol 2: 1350–1357.

Fuhrman, J.A. (1999) Marine viruses and their biogeochemical and ecological effects. Nature 399: 541–548.

Goldberg, S.M.D., Johnson, J., Busam, D., Feldblyum, T., Ferriera, S., Friedman, R., et al. (2006) A Sanger/pyrosequencing hybrid approach for the generation of high-quality draft assemblies of marine microbial genomes. Proc Natl Acad Sci U S A 103: 11240–11245.

Graham, E.D., Heidelberg, J.F., and Tully, B.J. (2017) BinSanity: unsupervised clustering of environmental microbial assemblies using coverage and affinity propagation. PeerJ 5: e3035.

Gutleben, J., Chaib De Mares, M., van Elsas, J.D., Smidt, H., Overmann, J., and Sipkema, D. (2018) The multi-omics promise in context: from sequence to microbial isolate. Crit Rev Microbiol 44: 212–229.

Handelsman, J., Tiedje, J., Alvarez-Cohen, L., Ashburner, M., Cann, I.K.O., Delong, E.F., et al. (2007) The New Science of Metagenomics : Revealing the Secrets of Our Microbial Planet, The National Academies Press.

Herrmann, M., Wegner, C.-E., Taubert, M., Geesink, P., Lehmann, K., Yan, L., et al. (2019) Predominance of Cand. Patescibacteria in Groundwater Is Caused by Their Preferential Mobilization From Soils and Flourishing Under Oligotrophic Conditions. Front Microbiol 10: 1407.

Howard-Varona, C., Hargreaves, K.R., Abedon, S.T., and Sullivan, M.B. (2017) Lysogeny in nature: mechanisms, impact and ecology of temperate phages. ISME J 11: 1511–1520.

Hug, L.A., Baker, B.J., Anantharaman, K., Brown, C.T., Probst, A.J., Castelle, C.J., et al. (2016) A new view of the tree of life. Nat Microbiol 1: 16048.

Hyatt, D., Chen, G.-L., LoCascio, P.F., Land, M.L., Larimer, F.W., and Hauser, L.J. (2010) Prodigal: prokaryotic gene recognition and translation initiation site identification. BMC Bioinformatics 11: 119.

Imachi, H., Nobu, M.K., Nakahara, N., Morono, Y., Ogawara, M., Takaki, Y., et al. (2019) Isolation of an archaeon at the prokaryote-eukaryote interface. bioRxiv.

Jain, C., Rodriguez-R, L.M., Phillippy, A.M., Konstantinidis, K.T., and Aluru, S. (2018) High throughput ANI analysis of 90K prokaryotic genomes reveals clear species boundaries. Nat Commun 9: 5114.

Kallies, R., Hölzer, M., Brizola Toscan, R., Nunes da Rocha, U., Anders, J., Marz, M., and Chatzinotas, A. (2019) Evaluation of Sequencing Library Preparation Protocols for Viral Metagenomic Analysis from Pristine Aquifer Groundwaters. Viruses 11.:

Kang, D.D., Froula, J., Egan, R., and Wang, Z. (2015) MetaBAT, an efficient tool for accurately reconstructing single genomes from complex microbial communities. PeerJ 3: e1165.

Kang, D., Li, F., Kirton, E.S., Thomas, A., Egan, R.S., An, H., and Wang, Z. (2019) MetaBAT 2: an adaptive binning algorithm for robust and efficient genome reconstruction from metagenome assemblies, PeerJ Preprints.

Kohlhepp, B., Lehmann, R., Seeber, P., Küsel, K., Trumbore, S.E., and Totsche, K.U. (2017) Aquifer configuration and geostructural links control the groundwater quality in thin-bedded carbonate–siliciclastic alternations of the Hainich CZE, central Germany. Hydrol Earth Syst Sci 21: 6091–6116.

Kurtz, S., Phillippy, A., Delcher, A.L., Smoot, M., Shumway, M., Antonescu, C., and Salzberg, S.L. (2004) Versatile and open software for comparing large genomes. Genome Biol 5: R12.

Küsel, K., Totsche, K.U., Trumbore, S.E., Lehmann, R., Steinhäuser, C., and Herrmann, M. (2016) How Deep Can Surface Signals Be Traced in the Critical Zone? Merging Biodiversity with Biogeochemistry Research in a Central German Muschelkalk Landscape. Front Earth Sci Chin 4: 32.

Lehmann, R. and Totsche, K.U. (2020) Multi-directional flow dynamics shape groundwater quality in sloping bedrock strata. J Hydrol 580: 124291.

Li, H. (2018) Minimap2: pairwise alignment for nucleotide sequences. Bioinformatics 34: 3094–3100.

Li, H., Handsaker, B., Wysoker, A., Fennell, T., Ruan, J., Homer, N., et al. (2009) The sequence alignment/map format and SAMtools. Bioinformatics 25: 2078–2079.

Luo, C., Tsementzi, D., Kyrpides, N.C., and Konstantinidis, K.T. (2012) Individual genome assembly from complex community short-read metagenomic datasets. ISME J 6: 898–901.

Matsen, F.A., Kodner, R.B., and Armbrust, E.V. (2010) pplacer: linear time maximum-likelihood and Bayesian phylogenetic placement of sequences onto a fixed reference tree. BMC Bioinformatics 11: 538.

Mikheenko, A., Saveliev, V., and Gurevich, A. (2016) MetaQUAST: evaluation of metagenome assemblies. Bioinformatics 32: 1088–1090.

Murat Eren, A., Esen, Ö.C., Quince, C., Vineis, J.H., Morrison, H.G., Sogin, M.L., and Delmont, T.O. (2015) Anvi’o: an advanced analysis and visualization platform for ‘omics data. PeerJ 3: e1319.

Nooij, S., Schmitz, D., Vennema, H., Kroneman, A., and Koopmans, M.P.G. (2018) Overview of Virus Metagenomic Classification Methods and Their Biological Applications. Front Microbiol 9: 749.

Nurk, S., Meleshko, D., Korobeynikov, A., and Pevzner, P.A. (2017) metaSPAdes: a new versatile metagenomic assembler. Genome Res 27: 824–834.

Parks, D.H., Chuvochina, M., Waite, D.W., Rinke, C., Skarshewski, A., Chaumeil, P.-A., and Hugenholtz, P. (2018) A standardized bacterial taxonomy based on genome phylogeny substantially revises the tree of life. Nat Biotechnol 36: 996–1004.

Parks, D.H., Imelfort, M., Skennerton, C.T., Hugenholtz, P., and Tyson, G.W. (2015) CheckM: assessing the quality of microbial genomes recovered from isolates, single cells, and metagenomes. Genome Res 25: 1043–1055.

Paulson, J.N., Stine, O.C., Bravo, H.C., and Pop, M. (2013) Differential abundance analysis for microbial marker-gene surveys. Nat Methods 10: 1200–1202.

Pedron, R., Esposito, A., Bianconi, I., Pasolli, E., Tett, A., Asnicar, F., et al. (2019) Genomic and metagenomic insights into the microbial community of a thermal spring. Microbiome 7: 8.

Price, M.N., Dehal, P.S., and Arkin, A.P. (2010) FastTree 2 – Approximately Maximum-Likelihood Trees for Large Alignments. PLoS One 5: e9490.

Quast, C., Pruesse, E., Yilmaz, P., Gerken, J., Schweer, T., Yarza, P., et al. (2012) The SILVA ribosomal RNA gene database project: improved data processing and web-based tools. Nucleic Acids Res 41: D590–D596.

R Core Team (2014) R: A Language and Environment for Statistical Computing.

Roux, S., Brum, J.R., Dutilh, B.E., Sunagawa, S., Duhaime, M.B., Loy, A., et al. (2016) Ecogenomics and potential biogeochemical impacts of globally abundant ocean viruses. Nature 537: 689–693.

Roux, S., Enault, F., Hurwitz, B.L., and Sullivan, M.B. (2015) VirSorter: mining viral signal from microbial genomic data. PeerJ 3: e985.

Schloss, P.D., Westcott, S.L., Ryabin, T., Hall, J.R., Hartmann, M., Hollister, E.B., et al. (2009) Introducing mothur: Open-source, platform-independent, community-supported software for describing and comparing microbial communities. Appl Environ Microbiol 75: 7537–7541.

Scholz, M., Lo, C.-C., and Chain, P.S.G. (2014) Improved assemblies using a source-agnostic pipeline for MetaGenomic Assembly by Merging (MeGAMerge) of contigs. Sci Rep 4: 6480.

Seemann, T. (2015) Barrnap.

Shaiber, A. and Eren, A.M. (2019) Composite Metagenome-Assembled Genomes Reduce the Quality of Public Genome Repositories. MBio 10.:

Taubert, M., Stöckel, S., Geesink, P., Girnus, S., Jehmlich, N., von Bergen, M., et al. (2018) Tracking active groundwater microbes with D2O labelling to understand their ecosystem function. Environ Microbiol 20: 369–384.

Tyson, G.W., Chapman, J., Hugenholtz, P., Allen, E.E., Ram, R.J., Richardson, P.M., et al. (2004) Community structure and metabolism through reconstruction of microbial genomes from the environment. Nature 428: 37–43.

Uritskiy, G.V., DiRuggiero, J., and Taylor, J. (2018) MetaWRAP-a flexible pipeline for genome-resolved metagenomic data analysis. Microbiome 6: 158.

Venter, J.C., Remington, K., Heidelberg, J.F., Halpern, A.L., Rusch, D., Eisen, J.A., et al. (2004) Environmental genome shotgun sequencing of the Sargasso Sea. Science 304: 66–74.

Wang, Q., Garrity, G.M., Tiedje, J.M., and Cole, J.R. (2007) Naïve Bayesian Classifier for Rapid Assignment of rRNA Sequences into the New Bacterial Taxonomy. Appl Environ Microbiol 73: 5261–5267.

Wegner, C.-E., Gaspar, M., Geesink, P., Herrmann, M., Marz, M., and Küsel, K. (2019) Biogeochemical Regimes in Shallow Aquifers Reflect the Metabolic Coupling of the Elements Nitrogen, Sulfur, and Carbon. Appl Environ Microbiol 85.:

Wickham, H. (2009) ggplot2: elegant graphics for data analysis, Springer New York.

Wickham, H., Averick, M., Bryan, J., Chang, W., McGowan, L., François, R., et al. (2019) Welcome to the Tidyverse. Journal of Open Source Software 4: 1686.

Woyke, T., Doud, D.F.R., and Eloe-Fadrosh, E.A. (2019) Genomes From Uncultivated Microorganisms. Encyclopedia of Microbiology, 4e.

Wu, Y.-W., Simmons, B.A., and Singer, S.W. (2016) MaxBin 2.0: an automated binning algorithm to recover genomes from multiple metagenomic datasets. Bioinformatics 32: 605–607.

Yan, L., Herrmann, M., Kampe, B., Lehmann, R., Totsche, K.U., and Küsel, K. (2019) Environmental selection shapes the formation of near-surface groundwater microbiomes. Water Res 170: 115341.

